# Stimulatory effect of monoacylglycerol lipase inhibitor MJN110 on locomotion and step kinematics demonstrated by high-precision 3D motion capture in mice

**DOI:** 10.1101/2023.06.25.546437

**Authors:** Bogna M. Ignatowska-Jankowska, Aysen Gurkan Ozer, Alexander Kuck, Micah J. Niphakis, Daisuke Ogasawara, Benjamin F. Cravatt, Marylka Y. Uusisaari

## Abstract

The neuromodulatory endocannabinoid system is a promising target for therapeutic interventions. One of the well-known behavioral effects of cannabinoid CB_1_ receptor activation with exogenous ligands such as THC is the inhibition of locomotor activity. However, the behavioral effects of endogenous cannabinoids are not understood. Enhancing endocannabinoid signaling offers an advantageous therapeutic strategy with limited cannabimimetic side effects, but their effects on motor function remain unclear. To reveal even the finest changes in motor function during voluntary locomotor tasks in mice, we adapted a high-speed, high-resolution marker-based motion capture, which so far has not been available in freely moving mice. Here we show that inhibition of distinct endocannabinoid metabolic pathways produces opposite effects on locomotor behavior that differ from those induced by exogenous cannabinoid receptor ligands. Selective upregulation of endocannabinoids 2-arachidonoylglycerol (2-AG) or N-arachidonoylethanolamine (AEA, anandamide) with inhibitors of their degradation (MJN110 and PF3845, respectively), produced bidirectional effects: MJN110 enhanced and PF3845 suppressed locomotor activity. Consistent differences in whole-body movement and precise step kinematics were found under distinct treatments, while analysis of locomotory episodes revealed invariant temporal microstructure, pointing towards motivational rather than motor-related mechanisms of action. The results show that the effects of manipulations of endocannabinoid system on locomotion are more diverse than previously assumed and result in distinct kinematic phenotypes.

## Introduction

Broad therapeutic potential of approaches targeting endocannabinoid signaling, as well as expanding medicinal and recreational use of exogenous cannabinoids^1–3^ call for a thorough investigation of endocannabinoid system modulation. The high expression of cannabinoid receptors in the cerebellum and basal ganglia^4–6^ suggests that precise assessment of subtle motor effects of compounds targeting the endocannabinoid system is necessary. The research on the effects of cannabinoids on precise movement in rodents is, however, limited ^7–11^ but cannabinoids have been shown to have effects on eyeblink ^12^ and kinematics of gait in humans^13^.

Currently, accurate kinematic tracking of rodents largely relies on restricting animals’ movements^14^ or on creating artificial environments (such as transparent floors or locomotion in narrow confines) that may distort behavior and limit the translational potential of the results. As the current translational crisis has highlighted the weakness of animal experimental designs^15–18^ their improvement is now urgently needed so that they would more accurately reflect the complexity of behavior in natural environments. In nature, it is rare for mice to explore flat, smooth surfaces such as those commonly used in the context of laboratory experiments. Instead, their natural behavior involves locomoting predominantly on uneven terrain with complex demands for body kinematics: they balance, climb, jump, and swim. Assessment of such 3D kinematics in freely behaving rodents in various environments requires the use of high-precision 3D motion capture.

Despite its precision, robustness and common use in humans^19–23^ use of marker-based 3D motion capture in small animals such as mice and rats has been limited by technical difficulties ^14,15,23–27^ and markerless animal tracking has become the main tool for assessing behavior (e.g. Deep Lab Cut, DeepEthogram, EthoLoop, MoSeq^28–34^. These approaches have recently experienced significant advances in flexibility and ease-of-use, but achieving precise 3D trajectory tracking remains challenging.^15,35,36^ Even in human studies, markerless techniques show only limited agreement with marker-based methods even under optimal conditions and typically generate mean errors of 10%.^21–25,35–42^. Importantly, further development of markerless techniques heavily depends on comparison with marker-based data^22,36,41^ which serves as the ground truth in most advanced human studies. As marker-based kinematic data is missing for rodents, improvement of markerless techniques is stifled. ^36,37,42^.

Reaching towards precise and accurate kinematic recordings of unrestricted mice moving freely in three dimensions, we modified a marker-based 3D motion capture (MoCap) system commonly used for human and other large animal tracking (Qualisys)^43–45^ to examine how mouse locomotor activity and kinematics were affected by modulation of the endocannabinoid system.

Here we aimed to assess whether the upregulation of local endocannabinoid signaling by selective inhibition of their degradation^46–49^ leads to behavioral outcomes different from those seen after systemic administration of exogenous ligands that bind to all available receptors in the body. To tease apart effects of inhibiting degradation of two endocannabinoids, 2-arachidonoylglycerol (2-AG) and N-arachidonoylethanolamine (anandamide, AEA), we examine the effects of selective inhibitors of their degradation in freely behaving mice tracked with the marker-based MoCap system.

The results reveal distinct effects for the two endocannabinoid-enhancing treatments, that are also different from those seen after exogenous cannabinoid agonist administration. Our observations suggest fundamentally different mechanisms of actions of distinct cannabinoids and call for revision of the current view of mostly inhibitory effects of cannabinoids on behavior. Here, we explicitly demonstrated the stimulatory effect of MAGL inhibition not only on general activity but also on fine kinematics. The work shows the strength of marker-based 3D MoCap as a powerful and sensitive tool for examining subtle differences in rodent behavior across spatial and temporal scales.

## Methods

### Experimental subjects and drug administration

We used adult male C57BL/6A (CLEA, Japan) mice (10-12 weeks old at the beginning of experiments) that were subjected to systemic intraperitoneal (i.p.) administration of drugs followed by behavioral assessment. To modulate endocannabinoid signaling we used the latest generation of pharmacological tools including inhibitors of FAAH and MAGL enzymes (PF3845 and MJN110, respectively, kindly provided by Dr. Benjamin Cravatt’s laboratory, Scripps Institute, USA). As a positive control we used potent agonist of CB_1_ and CB_*2*_ receptors CP55,940 (Tocris, UK). All doses were selected based on our previously published experiments^46,50^. The drugs were administered in a vehicle solution consisting of ethanol, colliphor and saline (1:1:18 ratio). Drugs were given at a volume of 10 μ*l/g* of body mass. Enzyme inhibitors will be administered 120 min before testing to ensure endocannabinoid accumulation, direct agonist 15-30 min before testing and antagonists 10 min before inhibitors or agonists.

### Permanent marker implantation on skin surface

A major obstacle in developing 3D motion capture for small animals has been lack of markers appropriate for rodents. Skin-glued markers are of limited use in unrestricted rodents, as they attract the animal’s attention and distort behavior. Moreover, markers made of soft materials will not survive the attention and require frequent replacements. We have developed a new method of attachment of permanent stainless-steel markers on a surface of skin of mice, that is fast, simple and does not disturb behavior. 11-14 days before the start of behavioral experiments mice undergo a procedure of marker implantation that enables high-resolution behavioral monitoring with infrared (IR) cameras. Under full isofluorane anesthesia (2-3%), 5 pairs of stainless-steel markers were implanted on the skin at landmark locations of the body (Fig. 1a,b, Suppl. Fig. 1 and 2.). Markers are <14 mm long, diameter of 0.9-1.27 mm (18-20 gauge) stainless steel rods ending with screw-on ball on both sides. The weight of all the markers combined does not exceed 10% of the animal body weight. The screw-on balls residing at the skin surface are covered with a strongly infrared (IR)-reflective coating enabling IR cameras to track the location of the markers and thereby position of the mouse body and limbs. To implant the markers, 2 small (<1.5 mm) holes are made in the skin with an 18 gauge (1.27 mm) sterile needle under full anesthesia. The needle is then replaced with the piercing that has been disinfected by 70 % ethanol for at least 20 min. Markers (Fig.1, Suppl. Fig. 1, 2, 3) were placed in the landmark locations of animals’ body: on the back, hips, knees, ankles and shoulder blades. These locations have been chosen to allow for precise calculation of body movements during locomotion, the posture of an animal as well as fine changes in body kinematics, such as tremor, miscoordination and subtle swaying. Implants do not cause noticeable discomfort to the animals, and they can wear them for over a year allowing multiple within-subject tests. The under-skin implants are tightly attached into the connective tissue and thus their movement is less confounded than top-of-skin markers by the loose rodent skin that obscures large ranges of joint movement.

**Figure 1.**
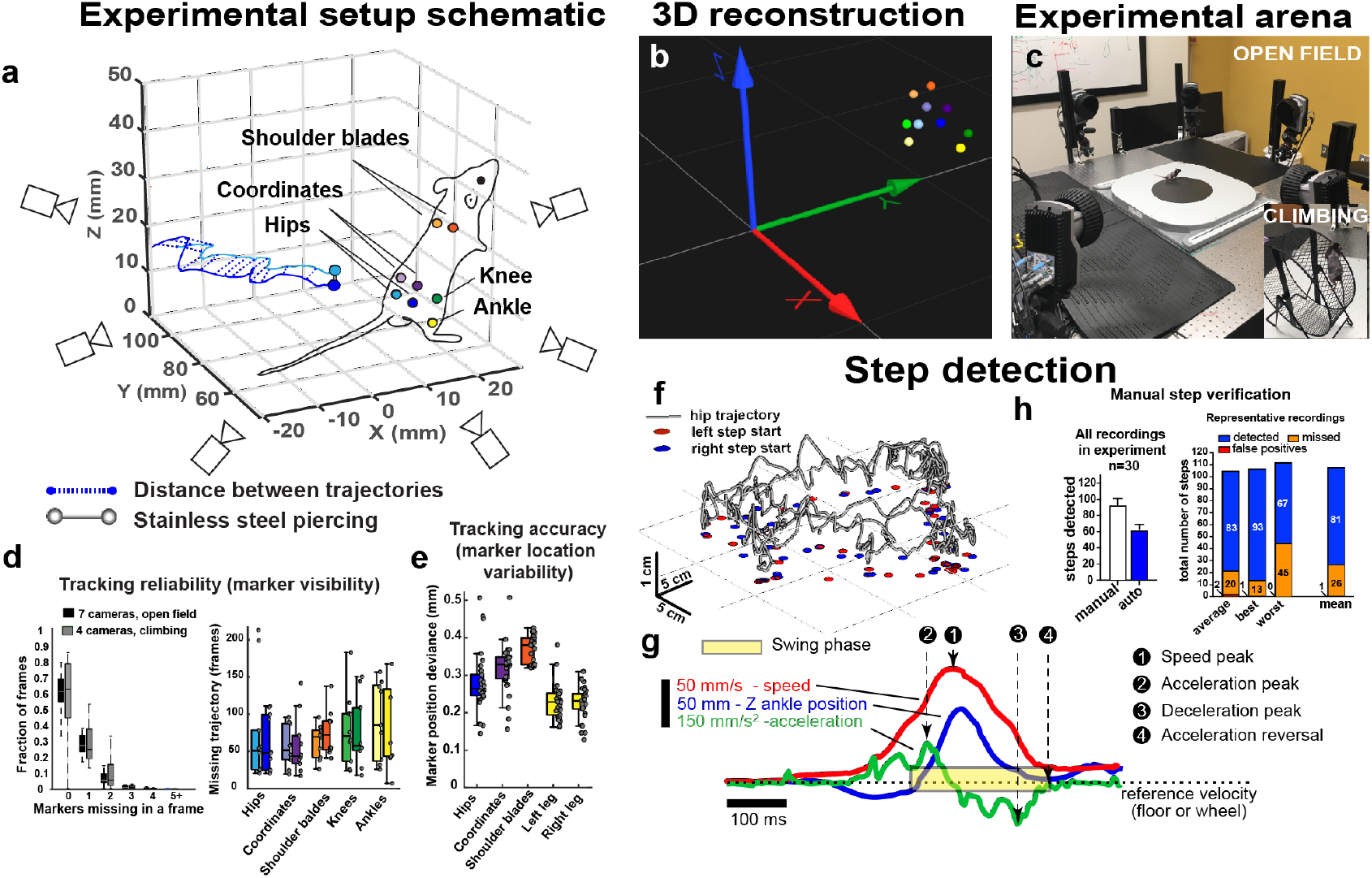
Motion Capture system adapted to mice allows for tracking movement trajectories with unprecedented precision. **a**. Schematic of experimental setup: mouse with retroreflective markers screwed on stainless steel implants (barbel piercings) at strategic locations on the skin surface, moving freely within 30 x 30 x 30 cm arena surrounded by 7 cameras. **b**. 3D reconstruction of marker locations using Qualisys motion capture system software. **c**. Photograph of the recording arena. **d**. Marker visibility determines tracking reliability. Left: fraction of frames in which markers are missing with 7 or 4 cameras (open field and climbing, respectively). Right: missing trajectory length for markers. Less than 100 frames are lost even in least visible markers. **e**. Marker pairs connected under the skin surface allow measurement of tracking accuracy based on marker location variability (distance between markers). Tracking accuracy falls between 0.15 mm and 0.5 mm for all markers (mean±SEM). **f**. Hip trajectories with steps marked during horizontal exploration by a representative animal. **g**. Step detection method. **i**. Manual step verification shows over 60% of steps are detected automatically with neglible number of false positives.

### Behavioral training

Spurious reflections and distortions caused by any transparent surfaces between the camera and markers catastrophically degrade tracking performance so during 3D recordings there can be no obstruction between cameras and an experimental subject, such as walls surrounding the experimental arena (Fig. 1 a-c). Therefore, animals require adequate training to a) habituate to naturally anxiogenic open field arena, b) train mice to remain in the arena during testing without any restriction of movement. We have developed a dedicated protocol for animal training allowing such behavioral assessment described in detail in supplementary methods. Experiments started 11-14 days following marker implantation to ensure proper skin healing and stable marker position. During those days animals were subjected to handling (3-5 days), acclimating to room and experimental arena (3-4 days), and task training: a) vertical climbing on mesh b) horizontal climbing on string c) beam walking d) righting reflex (2-3 days, 1 min trial or less). Markers that are worn by mice every day are exchanged into retroreflective and larger markers 15 minutes before the start of recording.

### Retroreflective marker preparation

Retroreflective markers are hand made by covering 4 mm diameter screw on stainless steel piercing ball (Fig. 1a, Suppl. Fig. 1a and 2) with highly retroreflective tape (3M, MN, USA). The tape is first cut into small pieces allowing covering ball with relatively smooth surface (Suppl. Fig. 3a). Then, tape is covered by layer of thin plastic (Linear Low Density Polyethylene (LLDPE) film, TRUSCO, Japan) followed by cover of liquid UV-cured plastic (Bondic, NY, USA) ensuring hard but transparent layer that will be sufficiently resistant to bites and other damage that may occur during testing.

### Assessing behavior using 3D motion capture

We adapted existing 3D motion capture system (Qualisys, Sweden) that is broadly used to study human and large animal biomechanics (Fig. 1a-c, Suppl. Video 1-5). Recordings were performed with 7 high-speed, high-resolution infrared-sensitive cameras (Qualisys Oqus 7, 12 megapixel sensor, 40 mm / f2.8 aperture) with integrated infrared-emitting LEDs, at 300 Hz allowing for highly precise recording and analysis of high-speed movement trajectories. Behavioral assessment was taking place within 30 x 30 x 30 cm in the center of the arena where cameras were focused. The open field arena is a 30 x 30 cm textured poliethylene floor (Suppl. Video 2,4). The climbing was performed on the outer surface of the metal mesh of the spoked running wheel for rats (25 cm in diameter). The wheel was moved manually according to voluntary mouse movement. Retroreflective markers were placed (spaced every 10 cm) on the outer side of the wheel to track it’s movement (Suppl. Video 3,5). All elements of the arena were cleaned in between trials.

### Calibrating motion capture recording for optimal tracking

To ensure reliable tracking of the retroreflective markers we calibrated the camera system at the beginning of each experimental day and every 4 hours during the day to avoid erratic tracking due to minute shifts in experimental system arrangements. We used the procedure provided by the hardware maker (Qualisys; https://docs.qualisys.com/getting-started/content/getting_started/running_your_qualisys_system/calibrating_your_system/calibrating_your_system.htm/) with slight modifications.

### Source code and kinematic feature extraction

Kinematic parameters were extracted by custom-made code in Matlab (2019a) run on MacOS Monterey 12.1. All code will be made available at GitHub repository at the time of publication and a link will be provided here. Methods of calculating key parameters computed from the 3D trajectories are described in supplementary methods.

### Performance of the marker-based mouse motion capture system

To ensure the reliability and accuracy of the measurements crucial in establishing a new method of recording behavior, we quantify them in terms of marker visibility (fraction of frames in a recording where a marker is visible or occluded) and location accuracy (how far is the recorded location from the real position of a marker). The vast majority of frames had no missing markers, and around 90% of frames had no more than one missing marker. The tracking quality was high even in climbing trials recorded with only 4 cameras (Fig. 1d, left). Most gaps lasted less than 100 frames (0.3 seconds at 300 fps) even in ankle markers that are naturally occluded in inactive, idle positions (Fig. 1d, right). The deviation of a marker’s tracked position relative to its pair can be considered as its error, and leg marker distance deviations of around 0.2 mm indicate that a tracked position of a single marker could be expected to be within 100 micrometers of its real position. The distance between paired markers varied on average no more than 0.5 mm (Fig. 1e). For step detection (Fig. 1f,g), we considered that a swing consists of a period of high-velocity ankle movement bordered by rapid acceleration and deceleration events. The number of steps detected by the algorithm was compared to those identified by an experienced observer, and the algorithm successfully detected majority of steps with less than 1% false positives (Fig. 1h, Suppl Fig. 3). More details are included in supplementary methods.

### Data analysis

Data are presented as mean ± standard error (SEM) and were analyzed using one-way or two-way analysis of variance (ANOVA). In case when some values were missing, these data were analyzed by fitting a mixed model, rather than by repeated measures ANOVA. Dunnett’s was used for post-hoc analysis in the dose–response experiments, and the Tukey or Sidak test was used for post-hoc analyses comparing different treatment groups. Differences were considered significant at the level of p<0.05. Statistical analysis was performed with GraphPad Prism version 8.00 (San Diego, CA) or Matlab (ver. 2019b, Mathworks, Natick, MA).

## Results

### 1. Validation of the 3D Motion Capture (MoCap) approach with well-known synthetic cannabinoid receptor agonist CP55,940

To validate the newly developed 3D MoCap system, we first chose to replicate previously-published results obtained with a well-known synthetic cannabinoid receptor agonist CP55,940^4,50^, before using compounds of less-known behavioral effects. We selected low doses (0.03, 0.1, 0.3 mg/kg), out of which only the highest dose was expected to result in any noticeable somatic effects in traditional assays ^50^ hypothesizing that the MoCap system might detect subtler behavioral changes than standard methods. The doses were verified in our laboratory with standard assay for triad of cannabimimetic effects (antinociception, hypothermia or catalepsy; Suppl. Fig. 4). The dose of 1 mg/kg was not included in motion capture experiments because it causes almost complete reduction of locomotor activity ^50^. The results were similar even though slightly smaller than previously observed^50^ indicating that the animals used in current study might be less sensitive to the doses than we observed in our earlier studies.

#### 1.1. General activity and locomotion parameters

We analyzed locomotor behavior in two behavioral tasks: horizontal exploration of the open field (Fig. 2a,b) and vertical climbing on a spoked wheel (Fig. 2c,d). We extracted measurements of changes in step kinematics as well as distance traveled, speed and general activity level, quantified as average of all markers’ speeds (“activity index”; Fig. 2a-d show example recordings for a representative individual treated either with vehicle or CP55,940 at 0.3 mg/kg). The results confirmed previous observations of a slight reduction of locomotor activity in the open field (Fig. 2b,e,f, Suppl. Fig. 5a-e, Suppl. Video 2). However, even with the highest dose used, trends towards decreased distance travelled (p = 0.36, Fig. 2f) and time spent locomoting (p = 0.18; Supp. Fig. 5c) were not significant and speed of locomotion (above 40 mm/s; p = 0.78, Suppl. Fig. 5d) was not affected. Interestingly, CP55,940 induced a significant dose-dependent decrease in the activity index (average speed of all markers) during 120 s recordings (F (3, 27) = 4.245, p = 0.014), which was most pronounced at 0.3 mg/kg (Fig. 2e,f), but also significant at 0.1 mg/kg (Suppl. Fig. 5a).

**Figure 2.**
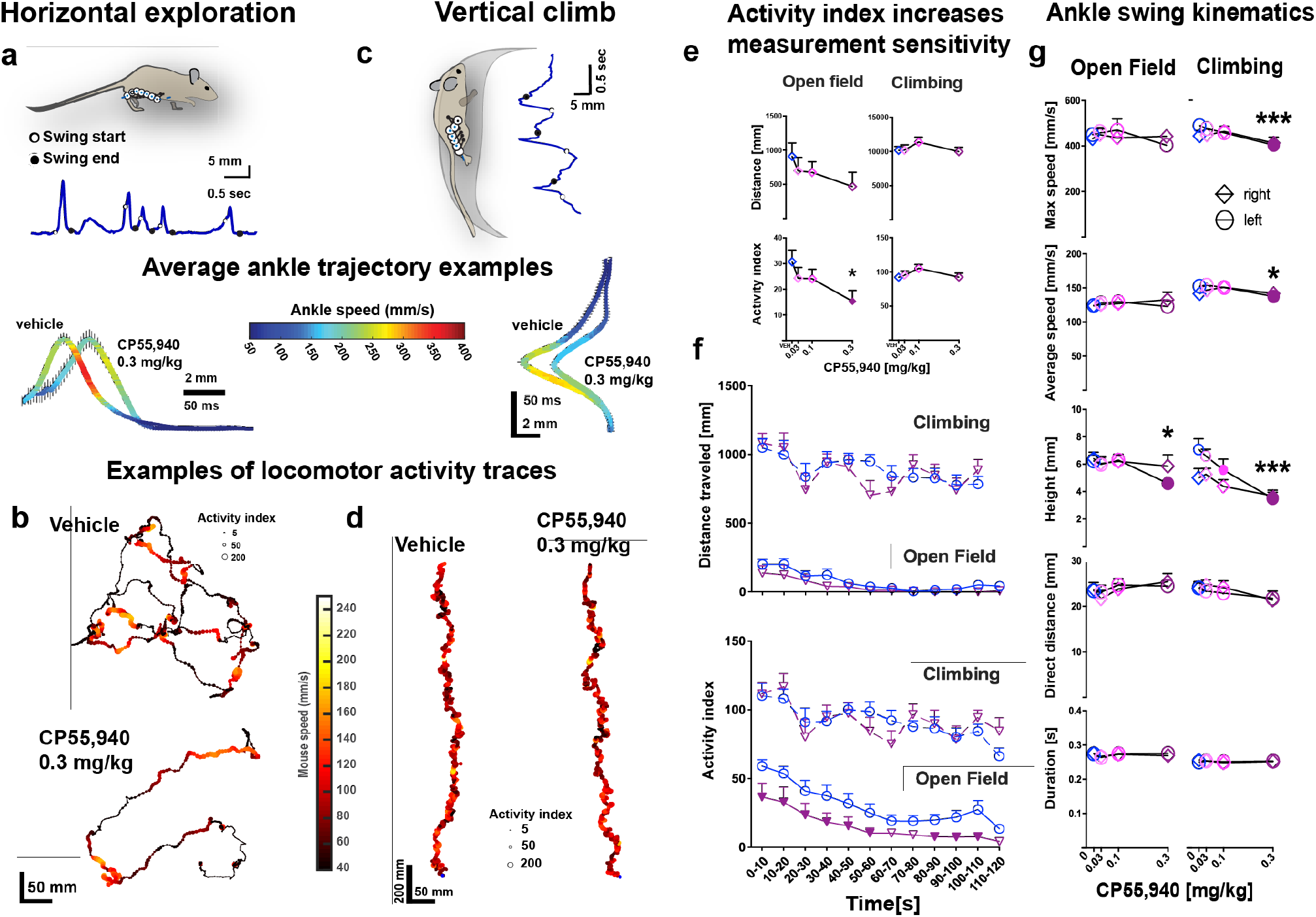
Marker-based 3D motion capture increases the sensitivity of behavioral analysis. **a**. Example of ankle trajectories during horizontal open field exploration: swing starts and stops (upper panel) and average ankle trajectory during all swings of representative animal following vehicle or CP55,940 (0.3 mg/kg) treatment. **b**. Example traces of locomotor activity in the open field following vehicle or CP55,940 at 0.3 mg/kg in representative animal. **c**. Example of ankle trajectories during vertical climbing: swing starts and stops (upper panel) and average ankle trajectory during swing of all steps of a representative animal following vehicle or CP55,940 (0.3 mg/kg) treatment. **d**. Example traces of locomotor activity during climbing following vehicle or CP55,940 at 0.3 mg/kg in representative animal. **e**. Total distance traveled (top) and mean activity index (bottom) during open field (left) and climbing (right). CP55,940 significantly decreased mean activity index at 0.3 mg/kg compared to vehicle, *p<0.05 (1-way ANOVA F (2.19, 19.75) = 4.25, Dunnett post hoc test) in the open field but not during climbing. **f**. Distance traveled (top) and activity index (bottom) as function of time during open field exploration and climbing. CP55,940 significantly decreased activity index during open field exploration. Closed symbols denote significance in Tukey post hoc test vs. vehicle (2-way ANOVA). g. CP55,940 significantly reduced maximum and average speed of ankle during climbing (right) and decreased step height in both open field (left) and climbing: *p<0.05, **p<0.01, ***p<0.001 post hoc comparison vs. vehicle (2-way ANOVA). e-g. n = 10 mice per group (randomized, counterbalanced within-subject design) mean±SEM.

To our surprise, no decrease in locomotor activity was observed in the climbing task, even though the mice performed it immediately after completing an open field task (Fig. 2d,e,f, Suppl. Fig 5f-j, Suppl. Video 3). In fact, a trend toward increased activity index following CP55,940 at 0.1 mg/kg during climbing could be discerned (Suppl. Fig. 5f).

A reduction of locomotor activity was observed towards the end of the trial in the climbing task (significant in all groups) but it was much less pronounced compared to the open field (Fig. 2a-f, Suppl. Fig. 5a-l).

There was no significant effect on stance width in either open field or climbing assays (data not shown), in line with the previous studies^4^ that reported widened stance only after a higher dose (1 mg/kg).

#### 1.2. Step kinematics parameters

The number of steps during trials was not significantly affected by CP55,940 in open field (p=0.27, Suppl. Fig. 5k) or climbing (p=0.23 Suppl. Fig. 5l). However, several step kinematic parameters were significantly altered (Fig. 2g, Suppl. Fig. 5m). Despite overall locomotion was not suppressed in the climbing task, significant slowing of ankle kinematics during swings was apparent with the highest dose (max ankle speed, F(3, 27) = 8.77, p<0.001; average ankle speed, F (3, 27) = 3.295, p = 0.036; Fig. 2g; Suppl. Fig. 5m shows similar measurements for knees). Moreover, CP55,940 significantly decreased height of ankle swing during climbing (F (3, 27) = 16.03, p<0.001), which was also observed in the open field (F (3, 27) = 4.474, p = 0.011); Fig. 2g) without other significant kinematic effects.

In summary, our results indicate that while CP55,940 inhibits exploratory locomotor behavior in the open field as previously reported^50^ even at moderate doses, it does not show such inhibition in the climbing task. Moreover CP55,940 at moderate doses significantly altered step kinematics by affecting the speed and height of the swing.

### 2. Effects of endocannabinoid signaling elevation on locomotor activity

Two best-known endogenous cannabinoids, 2-AG and AEA, are rapidly degraded by their respective catabolic enzymes, monoacylglycerol lipase (MAGL), and fatty acid amide hydrolase (FAAH) ^1,48,49^. To elevate concentrations of locally produced 2-AG and AEA we used selective inhibitors of MAGL and FAAH, MJN110 and PF3845, respectively. These inhibitors have been shown to reliably elevate 2-AG and AEA levels in the mouse central nervous system and peripheral tissues ^46,48,49^. To assess the behavioral effects of acute elevation of 2-AG and AEA in mice, we administered systemic MJN110 and PF3845 at relatively high doses that are effective in models of pain ^46,48,49^ and subjected mice to the same behavioral tasks as we used with CP55,940.

Inhibition of endocannabinoid degradation caused significant and opposite changes in locomotor activity and step kinematics during horizontal exploration of the open field and smaller but significant changes during vertical climbing (Fig. 3, Suppl. Fig. 6, Suppl. Video 4,5). Fig. 3a-d shows representative examples demonstrating changes in average ankle swing trajectories (Fig. 3a,c) as well as distance traveled, speed and general activity (quantified as activity index (AI); Fig. 2b,d) in a mouse treated with vehicle, PF3845 (30 mg/kg) and MJN110 (2.5 mg/kg). As compared in 2-way ANOVA, effects of PF3845 and MJN110 were bidirectional, where PF3845 caused significant inhibition of locomotor activity (Fig. 3a-f, Suppl. Fig. 6a-j) in all measures (distance traveled (F (2, 18) = 8.75, p = 0.022), activity index (F (2, 18) = 9.96, p = 0.012), time spent locomoting (F (2, 18) = 8.89, p = 0.021) and locomotion speed (F (2, 120) = 13.56, p<0.001)) compared to vehicle. In contrast, MJN110 induced significant increase in locomotor activity in the open field expressed as increase in distance traveled (Fig. 3a-f) and activity index (Fig. 3a-f) and increased time spent locomoting but had no significant effect on the speed as compared to vehicle. These bidirectional effects were also seen in climbing assay (Fig. 3d-f), as well as when lower doses were used (Suppl. Fig. 7). None of inhibitors affected stance width (data not shown).

**Figure 3.**
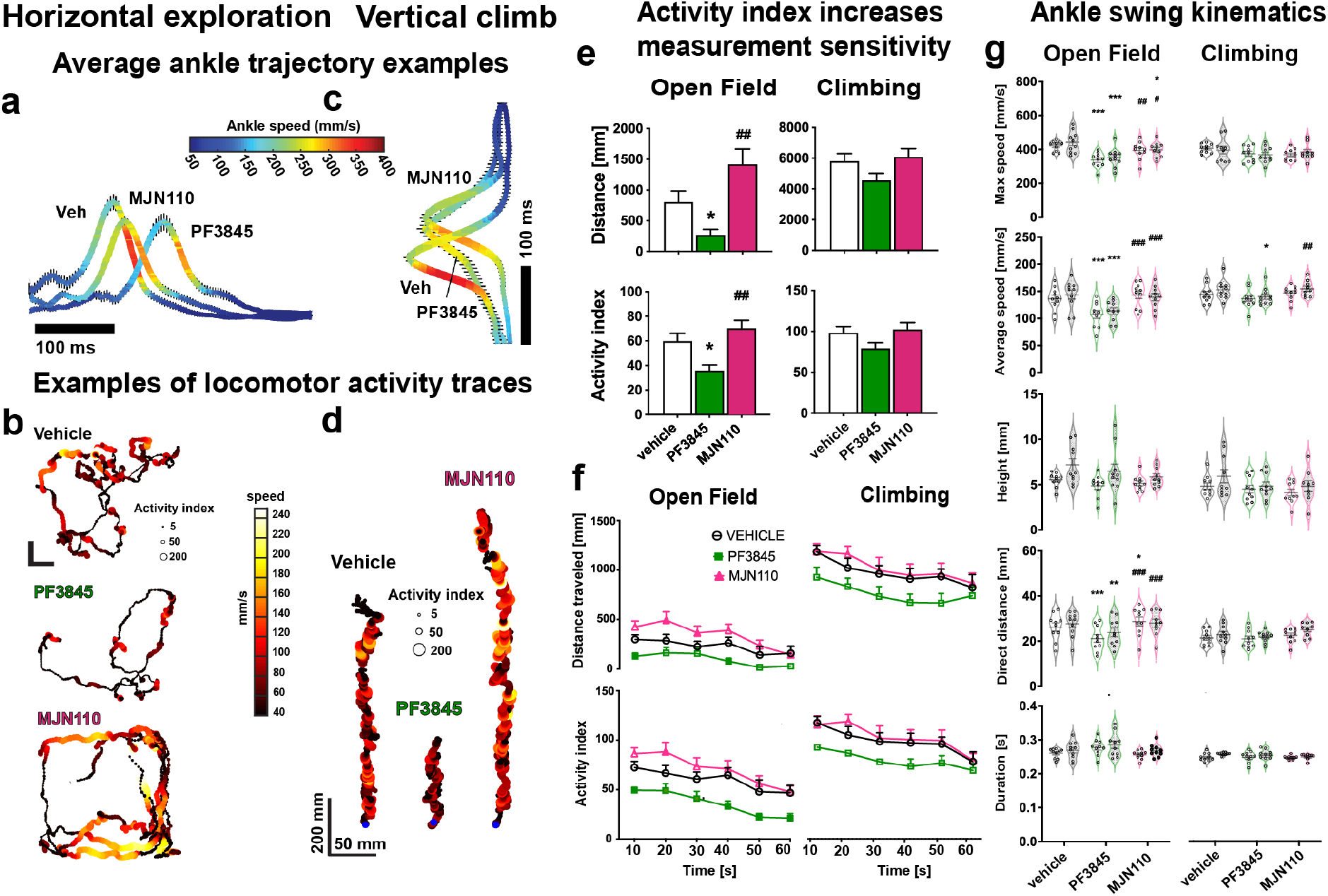
Opposite effects of distinct inhibitors of endocannabinoid degradation on locomotor activity and step kinematics. Examples of locomotor activity during open field exploration **(a, b)** and climbing **(c, d)** following treatment with PF3845 (30 mg/kg) or MJN110 (2.5 mg/kg) and compared to vehicle. **a** and **c:** Examples of average ankle trajectories SEM of all ankle swings of a representative animal during horizontal exploration and climbing, respectively. **b** and **d**: Examples of locomotor activity of a representative animal during horizontal exploration and climbing, respectively. **e**. Total distance traveled (top) and mean activity index (bottom) were significantly affected by treatment in horizontal exploration (left) (F (1.67, 15.06) = 10.52, F (1.92, 17.28) = 9.95), but not in climbing (right), *p<0.05, vs. vehicle; ##p<0.01 vs. PF3845 in post hoc comparison. **f**, left: Distance traveled (top) and activity index (bottom) were reduced (PF3845) and increased (MJN110) during the 60-s trials compared to vehicle control during horizontal exploration. **g**. PF3845 and MJN110 affected swing direct distances in bidirectional manner, so that PF3845 decreased them and MJN110 increased during open field exploration (left). Pairs of plots represent the right (right, shaded) and left leg (left, clear). The effects observed during climbing (right) were smaller than in open field. N=10 mice per group (randomized, counterbalanced within-subject design), mean±SEM.

### 3. Effects of endocannabinoid signaling elevation on step kinematics parameters

Similarly to effects on general locomotion, both inhibitors induced bidirectional effects on step kinematics. PF3845 (30 mg/kg) significantly decreased total number of ankle swings of both left and right leg during open field exploration (F (1.75, 15.73) = 10.45, p=0.002) and climbing task (F (1.79, 16.09) = 4.59, p=0.030) as compared to vehicle (Suppl. Fig. 6k,l). We have assessed differences in step kinematics parameters such as direct distance, total distance, height, duration, average speed, maximum speed in right and left leg under three treatment conditions: vehicle, PF3845, MJN110 (2-way ANOVA). PF3845 (30 mg/kg) significantly decreased direct (F (2, 18) = 3.75, p=0.043) and total distance (F (2, 18) = 4.83, p=0.021) as well as maximum (F (2, 18) = 24.09, p<0.0001) and average speed (F (2, 18) = 8.69, p=0.0023) of both right and left ankle (Fig. 3g). Similar effects of PF3845 were observed on swing of the knee (Suppl. Fig. 6m). MJN110 (2.5 mg/kg) significantly increased direct distance of the left ankle and knee swing compared to vehicle (Fig. 3g, Suppl. Fig. 6m). Less pronounced but significant effects on swing total distance and speed were observed in the climbing task (Fig. 3g, Suppl. Fig. 6m). There were no effects of treatment on duration or height of the swing of ankle or knee in open field or climbing (Fig. 3g). At lower doses (MJN110 1.25 and PF3845 10 mg/kg) we observed small but significant effects on step kinematics parameters of both compounds (Fig. 3g, Suppl. Fig. 7m).

### 4. Endocannabinoid modulation of locomotory episode structure

Bidirectional effects of inhibition of endocannabinoid degradation on locomotor activity were also reflected in the number of locomotory episodes defined by maintained displacement of the mouse at speeds over 40 mm/s. In the open field PF3845 induced significant decrease in total number of locomotory episodes as compared to vehicle (p = 0.013) and MJN110 (p=0.0026) (data not shown). In order to gain understanding of the mechanisms driving the bidirectional effects of MJN110 and PF3845 on locomotory activity, we analyzed the locomotory behavior of animals in more detail. Strikingly, we found that in all mice under all pharmacological conditions, the variable-duration locomotory episodes were composed of locomotory “bouts” during which the mice accelerate and decelerate in a stereotypical fashion. Comparing these bouts (shown for a representative example mouse in Fig. 4a1) under different drug conditions revealed that not only do the bouts maintain their speed profiles throughout the trials (as shown in Fig. 4a2 by comparing the bouts speed averages from first and last halves of the 60-s recordings), but also that the bouts are remarkably similar between different drug conditions (Fig. 4b1). Even in the PF3845 group, where the average locomotory speeds were significantly reduced in comparison to vehicle and MJN110 (Suppl. Fig. 6d), the duration of the bouts was virtually indistinguishable between the groups (bout half widths 1.190.003s, 1.360.006s and 1.22 s, ANOVA p = 0.06, Fig. 4b2). Indeed, the lower average speeds in PF3845 might result from reduced acceleration during the initial phase of locomotory bouts (quantified as the bouts slope; Fig. 4b3; pooled averages over entire trials 62.6 mm/s^*2*^, 42.3 mm/s^*2*^ and 65.1 mm/s^*2*^ for vehicle, PF3845 and MJN110; vehicle vs. PF3845, p < 0.0001; vehicle vs. MJN110, p = 0.46).

**Figure 4.**
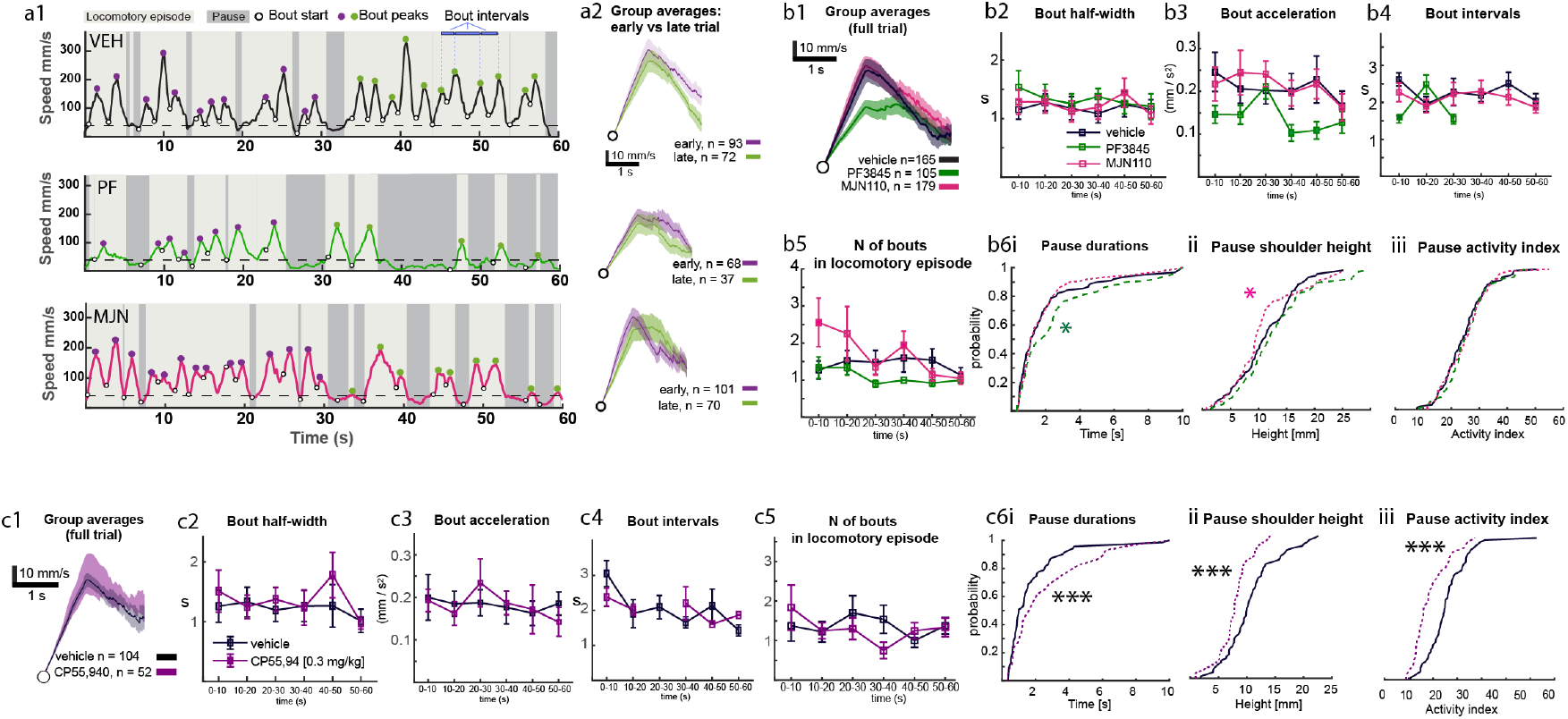
Invariant microstructure of locomotory episodes under modulation of endocannabinoid system. A, speed measurements during full 60-s trials in a single mouse in vehicle, PF3845 and MJN110 conditions. A1, speed during full trials; light and dark grey boxes denote locomotory episodes and pauses, respectively. Filled circles: speeds peaks (purple, first half of the trial; green, last half of the trial). Open circles: initiations of locomotory bouts. Dashed line indicates threshold for locomotion. A2, average SEM speeds in all locomotor bouts in all animals, divided into first and last-half events, aligned vertically to bout initiation (open circles). B, MJN110 does not cause changes in the shape of bouts (b1-3) or increase in likelihood of initiating locomotory episodes (b4) but increased number of bouts within the early episodes of locomotion (b5). PF3845 reduces the acceleration during a bout (b1, 3). Pause durations (b6i) and behavior during pauses (b6ii, iii) were not significantly affected. C, CP55,940 (0.3 mg/kg dose) did not cause changes in locomotory bout shape (c1-c3) nor their intervals or number within a locomotory episode (c4-c5). Instead, CP55,940 increased duration of locomotory pause durations (c6i), simultaneously lowering mouse posture and activity index (c6ii, iii). Closed symbols in panels b2-b5 denote significance in post hoc test vs. vehicle. Asterisk denote significant difference in distributions with K-S test vs. vehicle for the group indicated by color (b6) or ANOVA (c6) p* < 0.05, p***, < 0.001. n=10 mice per group, mean±SEM.

Another property of the locomotory episode structure clearly evident in the trial speed profiles (Fig. 4a1) was the invariance of bouts intervals; indeed, throughout the entire trials and in all pharmacological conditions, sequential bouts occurred at remarkably constant intervals (mean inter-bout intervals, 2.2, 2.1 and 2.1 s for vehicle, PF3845 and MJN110; ANOVA p >0.99, Fig. 4b4) throughout the locomotory episodes. Thus, the changes in time spent locomoting (Suppl. Fig. 6c) are reflected in the larger number of bouts during the early parts of the trials under MJN110 (1.42, 1.14 1.7 bouts per locomotory episode; 116, 92 and 105 episodes for vehicle, PF3845 and MJN110 groups, respectively; Fig. 4b5). In other words, the larger distance covered in MJN110 mice is due to them initiating more of these stereotypical bouts before pausing, while the mice under PF3845 initiate slightly fewer of these 1-2 -second-long bouts.

The fact that MJN110 influences the stopping likelihood of a locomoting mouse prompted us to ask whether it would also modulate the eagerness of the mice to initiate a locomotory episode. If this were the case, it should manifest as shortened duration of the pauses between locomotory episodes. However, as shown in Fig. 4b6i, the distribution of pause durations was nearly identical between vehicle and MJN110 (2.1 and 1.9 sek, K-S test 0.9), whereas the pauses were slightly longer in PF3845 (2.6 s; K-S 0.047 for PF3845 vs. vehicle). Thus, the increased exploratory behavior in MJN110 involves changes in mechanisms that control stopping rather than starting locomotion. The slightly longer pause durations caused by PF3845 do not suffice to explain the dramatic decrease in the distance the mice covered during PF3845 trials. Therefore we wondered if PF3845 would generally decrease muscle tone, leading to weaker acceleration and thereby eventual failure of reaching sufficient momentum for maintaining locomotion during the locomotory bout time windows. To investigate this, we examined the posture and activity measures during the locomotory pauses (Fig. 4b6ii-iii). Somewhat surprisingly, no dramatic differences were found, despite slight lowering of the shoulder height in MJN110 could be seen (11.2, 12.7, 10.6 mm; ANOVA p = 0.062, K-S PF3845 vs. MJN110 p = 0.005). These results obtained from quantification of the invariant locomotory bouts suggest that the bidirectional effects of MJN110 and PF3845 are mainly realized by modulation of motivational parameters of behavior: PF3845 makes the mice slower in initiating a locomotory bout. In contrast, MJN110 upregulates the likelihood that a locomoting mouse will initiate another bout instead of stopping, leading to overall longer distance covered.

These results with the selective endocannabinoid degradation enzyme inhibitors are in stark contrast with those obtained with the exogenous cannabinoid receptor agonist CP55,940 (Fig. 4c). While the bout shapes and intervals were virtually identical in mice under the influence of CP55,940 (panels c1 – c4; bout half-widths 1.25, and 1.39 sek; bout accelerations 54.9 and 55.3, p = 0.93; bout intervals 2.0 and 2.1 s for vehicle and 0.3 mg/kg CP55,940), the decrease of distances mice travelled was not caused by shortening of locomotory episode duration or number of bouts per episode (1.35 and 1.41 bouts in an episode in vehicle vs. CP55,940; n = 77 and 37 locomotory episodes; p >0.99; Fig. 4c5). Instead, nonspecific activation of cannabinoid receptors with CP55,940 increased duration of pauses between locomotory episodes (Fig. 4c6i; 1.73 vs. 2.6 s; ANOVA p < 0.001; n = 71 and 29 pauses in the first 60 s of the trials), concomitantly with lowering of overall posture (shoulder height; Fig. 4c6ii, 10.8 vs. 8.0 mm; ANOVA p < 0.001) and activity index (Fig. 4c6iii, 25.6 vs. 19.0, ANOVA p < 0.001) during the pauses.

Taken together, even though selective FAAH inhibition with PF3845 affected behavior in a way that superficially resembles that of activation of CB_1_ receptor with exogenous agonist CP55,940, the underlying features are different and likely involve different targets.

## Discussion

Taking advantage of the high precision and accuracy of our markerless 3D MoCap system, we revealed that selective inhibition of the degradation of endocannabinoids 2-AG and AEA results in distinct, bidirectional kinematic phenotypes that markedly differ from the effects of an exogenous synthetic ligand of the cannabinoid receptor. To our surprise, we also found that the effects were not universal to locomotory activity but differed between horizontal and vertical locomotion.

The endocannabinoid system, consisting of endogenous cannabinoids, their receptors and enzymes responsible for their biosynthesis and degradation^1^ is of high interest in the development of treatments for various diseases including neurodegenerative, autoimmune, and pain-related disorders ^1–3,51,52^. The medicinal compounds targeting cannabinoid receptors have relatively mild side effects (e.g. subjective intoxication, inhibition of locomotion and memory impairment^53^ and future studies aiming at the reduction of the cannabimimetic side effects will further increase their attractiveness for widespread therapeutic use^1,46,47,50^. Thus in the present study we did not aim to evoke dramatic behavioral distortions but focused on more subtle effects that might be indicative of distinct interoceptive states, difficult to observe with traditional behavioral assessment methods ^46,47^

The profound locomotion-inhibiting effect of the CB_1_ receptor activation by high doses of exogenous compounds is well known^4,54^. However, even though three decades have passed since the initial characterization of endogenous cannabinoids ^3,55^, their rapid degradation in living animals has limited research into their roles in behavior. Moreover, effects of synthetic and phyto-cannabinoid (e.g. delta-9-THC) on movement have mostly been examined using coarse behavioral assays as well as high doses unsuitable for widespread therapeutic use.

Recently-developed potent and selective inhibitors of the enzymes responsible for endocannabinoid degradation can be used to reveal effects of enhancing endocannabnoid signaling produced naturally *in situ* ^46–48,56^ unlike direct exogenous compounds targeting the CB_1_ receptor. We and others have shown that administration of these highly selective FAAH and MAGL inhibitors have effects similar to direct ligands, suggestive of therapeutic potential with less cannabimimetic side effects^46– 49.56–59^

According to ours and others’ unpublished observations various MAGL inhibitors ^46,47^ produce phenotype characterized by jittery rapid movements. Somewhat similar hyperreflexia manifested as jumping and “popcorning” behavior is observed after administration of large doses of cannabinoid agonists.^4,54^ However, at such doses drastic decline in spontaneous locomotion as well as other cannabimimetic effects are observed. It must be noted that MAGL inhibitors with very low cross-reactivity with FAAH (such as MJN110 and KML29) are known to not inhibit locomotion unlike JZL184 that has significant cross-reactivity with FAAH^46,47,60^. Highly selective MAGL inhibitors seem to show some stimulatory effects on activity^46,47^, but the nature of the phenotype their induce remains to be evaluated.

Endocannabinoids interact with multiple receptors besides CB_1_ and CB_*2*_ and manipulating their availability is expected to produce different effects than compounds such as CP55,940, designed to target CB_1_ and CB_2_ receptors^1^. As the ligand-receptor interactions differ in a cell- and tissue-type-dependent manner ^61–63^ and expression of receptors and enzymes is highly diverse across animal bodies^1^, it should not, in fact, be a surprise that each of the treatments in this study led to a distinct behavioral phenotype. More research is necessary to establish mechanisms underlying observed differences and establish molecular targets involved, which was beyond the scope of this article.

In the context of therapeutic use considerations, it is interesting to note that the selective MAGL inhibitor MJN110, expected to enhance 2-AG signaling, led to a clear stimulatory effects on locomotion, including increase in hindlimb swing distance (Fig. 3, Suppl. Fig. 6). This is in line with our and others’ anectodal observations of mild stimulatory effects of MAGL inhibition^46,64^. Moreover, most recent studies suggest that elevated 2-AG signaling might be associated with hyperdopaminergic and propsychotic outcomes ^65–67^. Considering the known reduction of locomotion and emergence of suicidal ideation caused by CB_1_ receptor antagonism^68,69^, it seems possible that 2-AG, the most abundant endogenous cannabinoid in the central nervous system, would play a stimulatory or pro-dopaminergic role in animal behavior and suppressing its signaling would lead to the negative symptoms.

In contrast, enhancing AEA signaling with inhibition of FAAH resulted in locomotory suppression, and the effects were consistent in the behavioral tasks. However, while the cannabinoid receptor agonist CP55,940 with known behavior-suppressing effects did reduce locomotion in horizontal exploration, no such inhibition was seen in vertical climbing task (Fig. 2). This observation clearly indicates that inhibition of exploration in the open field is not a result of impaired locomotory coordination and suggests that the suppression may consist of a significant motivational component, possibly combined with effect on muscle tone. Furthermore, it highlights the importance of using multiple behavioral tasks in order not to miss effects that might be context-dependent.

In addition to observing the effects on general locomotory behavior that could be in principle assessed with simple whole-animal tracking, the marker-based MoCap recordings revealed several more subtle signatures of animal behavior. First, we note that the magnitude of animal movement (quantified as “activity index”; Fig. 2-3) is more sensitive measurement of behavioral suppression or enhancement than traditional measures such as distance traveled. Second, the bidirectional effects on general locomotor activity were accompanied by significant changes in step kinematics, that were also bidirectional (Fig. 3). While PF3845-treated mice generated less steps that were shorter in distance and slower, MJN110-treated mice made steps that were longer. Interestingly, across all experiments the most pronounced changes in step kinematics parameters were observed in swing speeds, some changes were observed in swing distance or height, but step durations were remarkably unchanged. Third, a comparison of behavioral microstructure features (hindpaw swing duration, figures 2-3, and “bouts” of speed within locomotory episodes, Fig. 4) revealed surprising invariance: across all animals, all groups and throughout the trials, durations of swings and locomotory bouts were virtually identical in both horizontal and vertical locomotion. Thus, the CP55,940 and PF3845 did not make mice cover shorter distances by shortening their steps or locomotory “bouts” durations, but by decreasing the acceleration when initiating those events. Conversely, the striking enhancement of locomotory activity induced by MJN110 was not accompanied by any changes in locomotory “bout” in comparison to vehicle-treated condition, but by elevated likelihood of initiating more bouts within a locomotory episode before pausing the explorative activity. Finally, comparing animal posture and behavior during the pauses between locomotory episodes suggests that while direct activation of CB_1_ receptors with a low dose of CP55,940 and enhancement of AEA-mediated endocannabinoid signaling similarly suppressed locomotion, the latter is associated with less cannabimimetic side effects even at high doses.

Due to limitations caused by the low experimental throughput of our system while we were developing it, we needed to restrict our study to only one sex. The decision to use male mice in the first study using this methodology in mice was driven by the need to compare our results to earlier studies conducted only in male mice. It is necessary to establish differences in fine kinematics between male and female mice, and it is likely that observed effects of endocannabinoid system modulation could be different in female mice.

Our marker-based mouse motion capture approach opens new possibilities for studying the intricate kinematics of mouse movement by eliminating the need for using machine learning methodologies in feature extraction. This allows employing advanced mathematical and information-theoretical approaches to investigate the neurobiological basis of behavior in a more direct and powerful manner than previously. In the scope of rodent behavioral neuroscience, marker-based motion capture offers a significant advantage over classical and markerless methods due to its superior precision and extensive range in both spatial and temporal domains. This enables simultaneous analysis of behaviors ranging from speed profiles of individual steps with a resolution of tens of milliseconds, to whole-body movement trajectories spanning minutes or longer.

While further research into the systemic, cellular and molecular components underlying these effects is needed, the work presented demonstrates not only that marker-based 3D motion capture is an attainable method to apply in mice and other laboratory rodents, but also that it allows investigation of small movements (such as tremor; unpublished observations by the authors) in entirely unrestricted animals. Such spatiotemporal precision may reveal novel, subtle kinematic symptoms of degenerative diseases and support targeted development and refinement of interventions. Finally, the simplicity and precision of the method, combined with analysis that does not depend on machine learning or model-fitting approaches opens doors for more creative experimental design involving natural behaviors, such as hunting and social interactions, and could identify kinematic signatures of interoceptive states that currently are extremely challenging to define in rodent models ^46^.

## Data availability

All motion capture tracking files including labeled data will be fully available for download at the time of publication.

## Authors’ contributions

BIJ: experimental design, conducting experiments, data analysis, manuscript writing, visualization

AK: conducting experiments, programming, manuscript revision

AGO: conducting experiments, manuscript revision

DO: drug design and synthesis, manuscript revision

MJN: drug design and synthesis, manuscript revision

BFC: drug design and synthesis, manuscript revision

MYU: software design and programming, data analysis, manuscript writing, visualization

All authors have read and agreed to the published version of the manuscript.

## Ethics approval

The animal study protocol was conducted in accordance with procedures approved by the Okinawa institute of Science and Technology (OIST) Institutional Animal Care and Use Committee (IACUC) (Protocol IDs: 2017-188, 2020-305) in accordance with the National Institutes of Health Guide for the Care and Use of Laboratory Animals (National Research Council, 2011) Every effort was made to minimize suffering.

