## Supplementary Materials for "Stimulatory effect of monoacylglycerol lipase inhibitor MJN110 on locomotion and step kinematics demonstrated by high-precision 3D motion capture in mice"

*3D MoCap reveals distinct kinematic phenotypes of endocannabinoids in mice, Ignatowska-Jankowska et al.*

### <u>Supplementary Figures</u>

#### Supplementary Figure 1.

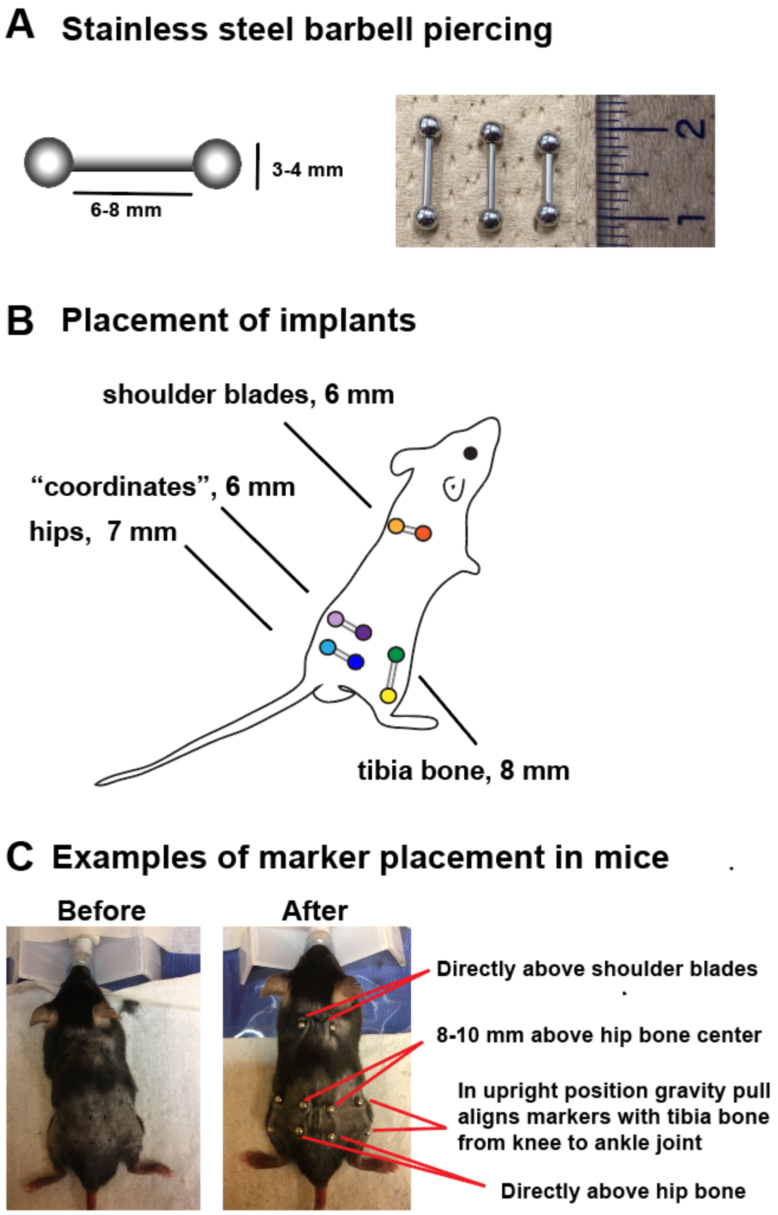


**Supplementary Figure 1. Implantation of markers a,** Stainless steel implants (barbel piercings) used to attach retroreflective markers during experiments. **b,** Localization of marker placement on the mouse skin. **c,** Example of placement of the markers on representative mouse before and after the implantation.

### Supplementary Figure 2

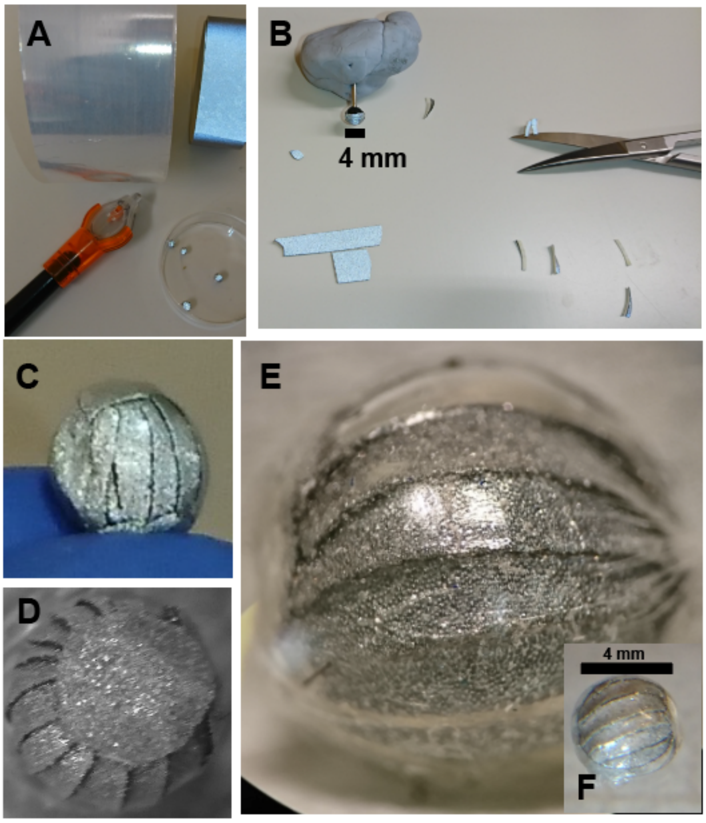


**Supplementary Figure 2. Retroreflective marker preparation. a,** Materials for marker preparation: retroreflective tape (top right), polyethylene film (top left), UV cured Bondic glue (bottom left). **b,** Thin strips of retroleflective tape glued on 4 mm stainless steel ball. **c,** Steel ball covered with cut retroreflective tape. **d,** Steel ball covered with tape and poliethylene film. **e,f,** Finished marker: steel ball covered with tape, polyethylene film and UV hardened Bondic glue.

### Supplementary Figure 3

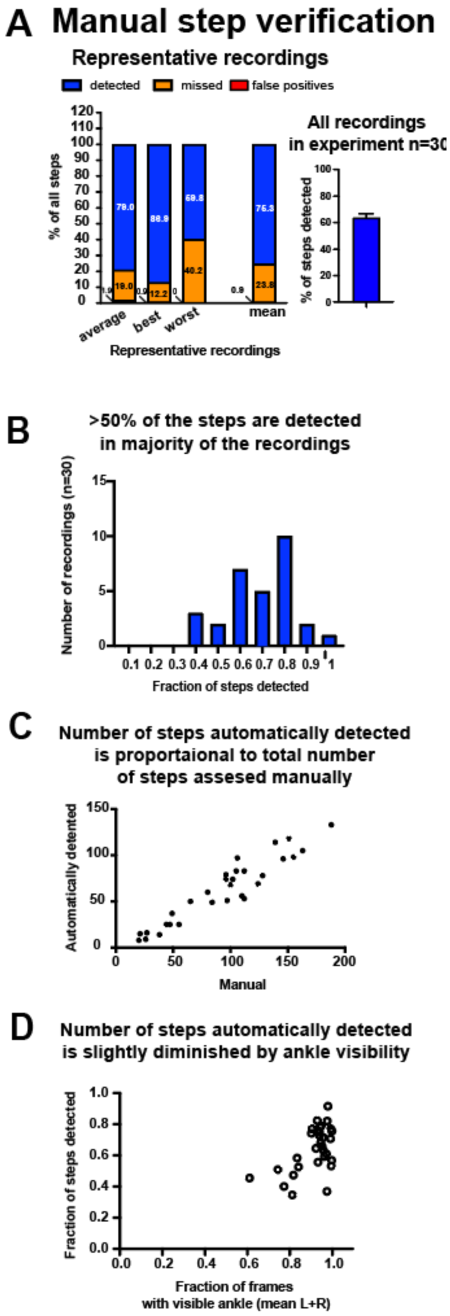


**Supplementary Figure 3. Manual step detection verification**
**a,** Percentage of steps detected in representative recordings (left panel) as well as in total of 30 recordings in an experiment (right panel). Step detection was optimized to minimize false positives while preserving sufficient number of steps recorded for analysis. **b,** Histogram showing number of recordings with fraction of steps detected. **c,** Automatic step detection shows similar results to manual verification. **d,** While visibility of markers can disturb step detection, in our experiments step detection was only mildly affected by ankle visibility.

### Supplementary Figure 4

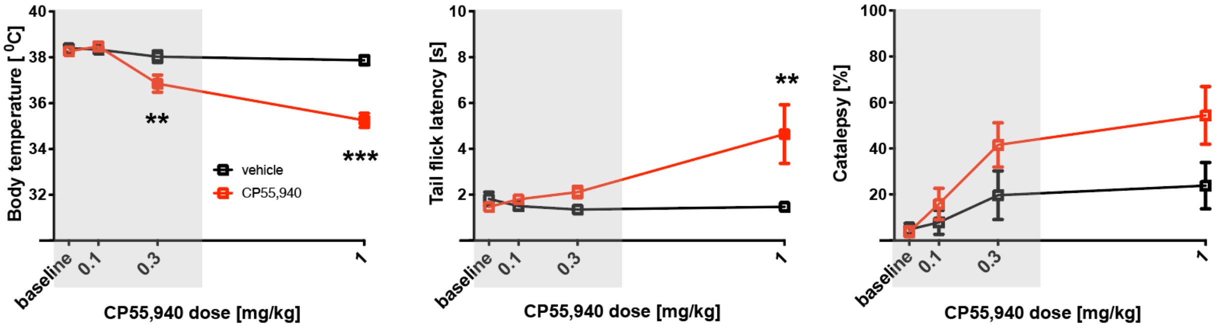


**High dose of CP55,940 (1 mg/kg) produces cannabimimetic effects: hypothermia, analgesia and catalepsy in cumulative triad assay, while moderate doses induce minimal effects.** CP55,940 treatment caused**:** **a,** significantly reduced body temperature at 0.3 and 1 mg/kg ($F (1, 55) = 31.08$, $p<0.0001$), **b,** significant increase antinociception in tail flick test at 1 mg/kg ($F (1, 55) = 7.809$, $p=0.007$), and **c,** significantly increased catalepsy (main effect of treatment $F (1, 63) = 6.091$, $p=0.016$). 2-way ANOVA, followed by Sidak post hoc comparisom, **p<0.01, ***p<0.001 vs vehicle treatment. Mean±SEM, n=10 per group.

### Supplementary Figure 5

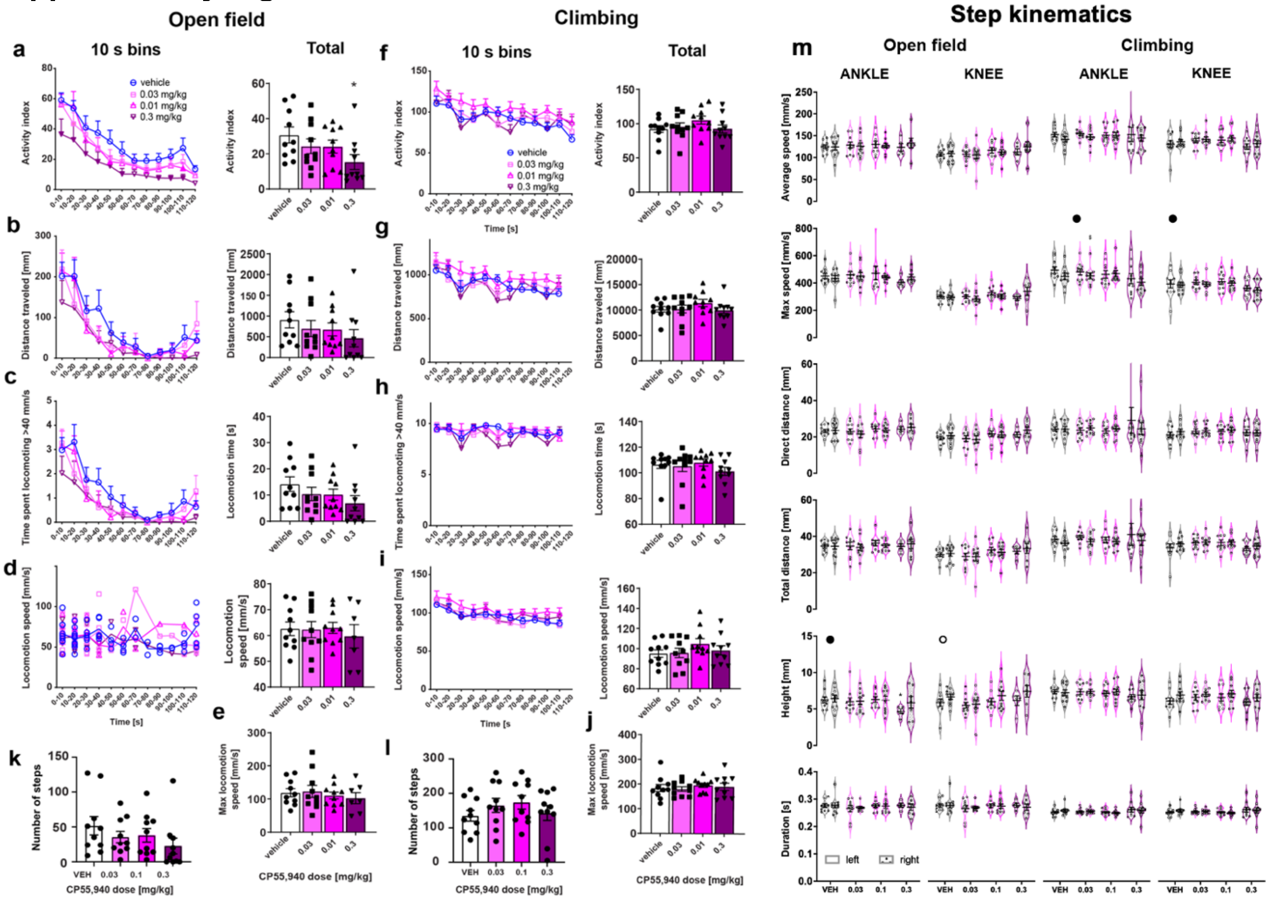


**Supplementary Figure 5. Low dose CP55,940 decreases activity in the open field but not in the vertical climbing task.**

**a,** CP55,940 induced dose-dependent, significant decrease in activity index (average speed of all markers) during exploration of the open field. Closed symbols denote significance in Dunnett post hoc test vs vehicle (left, 2-way ANOVA), *p<0.05 vs vehicle (right, 1-way ANOVA). **b,** Distance traveled and **c**, time spent locomoting (above 40 mm/h) was not significantly affected. **d**. No change in average or **e.** maximum locomotion speed was observed. During climbing CP55,940 did not have any significant effects on **f**, activity index, **g,** distance traveled, **h**, time spent locomoting or **i,** average or **j,** maximum locomotion speed during. **k,** CP55,940 did not affect number of steps during open field or **l,** climbing trials. **m,** CP55,940 had small but significant effects on step kinematics on maximum and average speed as well as swing height. Black circles indicate significant treatment effects in 2-way ANOVA. Open circle denotes significant effect of leg side. n=10 mice per group (randomized, counterbalanced within-subject design), mean ± SEM.

### Supplementary Figure 6

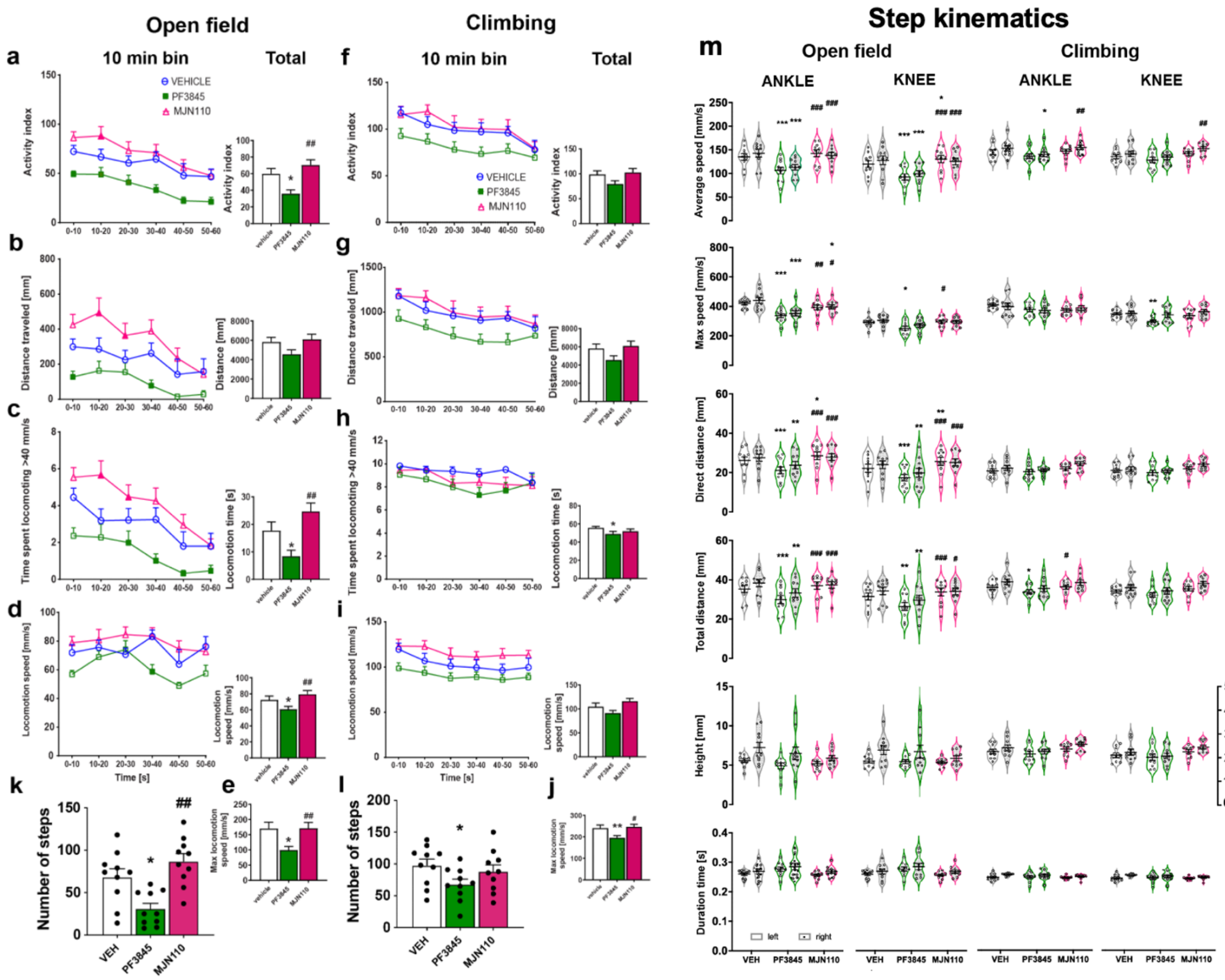


**Supplementary Figure 6. MAGL and FAAH inhibitors PF3845 (30 mg/kg) and MJN110 (2.5 mg/kg) produce bidirectional effects on locomotor activity and step kinematics.** In the open field, PF3845 significantly decreased and MJN110 significantly increased: **a,** activity index, **b,** distance traveled **c,** time spent locomoting, compared to vehicle. **d,** PF3845 significantly decreased average, and **e,** maximum locomotion speed, while MJN110 had no significant effect on speed compared to vehicle. **f,** During climbing both inhibitors had similar but less pronounced effects: **g,** activity index, and **h,** distance traveled were decreased by PF3845 but effects were not statistically significant (p=0.068, p=0.064). **h,** PF3845 induced significant decrease in locomotion time during climbing (F (2, 18) = 3.79, p<0.042). **i,** Significant effect of treatment was found on locomotion speed (F (2, 162) = 19.04, p<0.001) during climbing. **k,** PF3845 significantly decreased number of steps during open field and also **l,** during climbing trial. **m,** Similar bidirectional effects on swing kinematics were also found on swing distance, but not height or duration. *p<0.05, **p<0.01 vs vehicle, #p<0.05 ##p<0.01 vs PF3845. Closed symbols denote significance in post hoc test vs vehicle. n=10 mice per group (randomized, counterbalanced within-subject design), mean ± SEM.

### Supplementary Figure 7

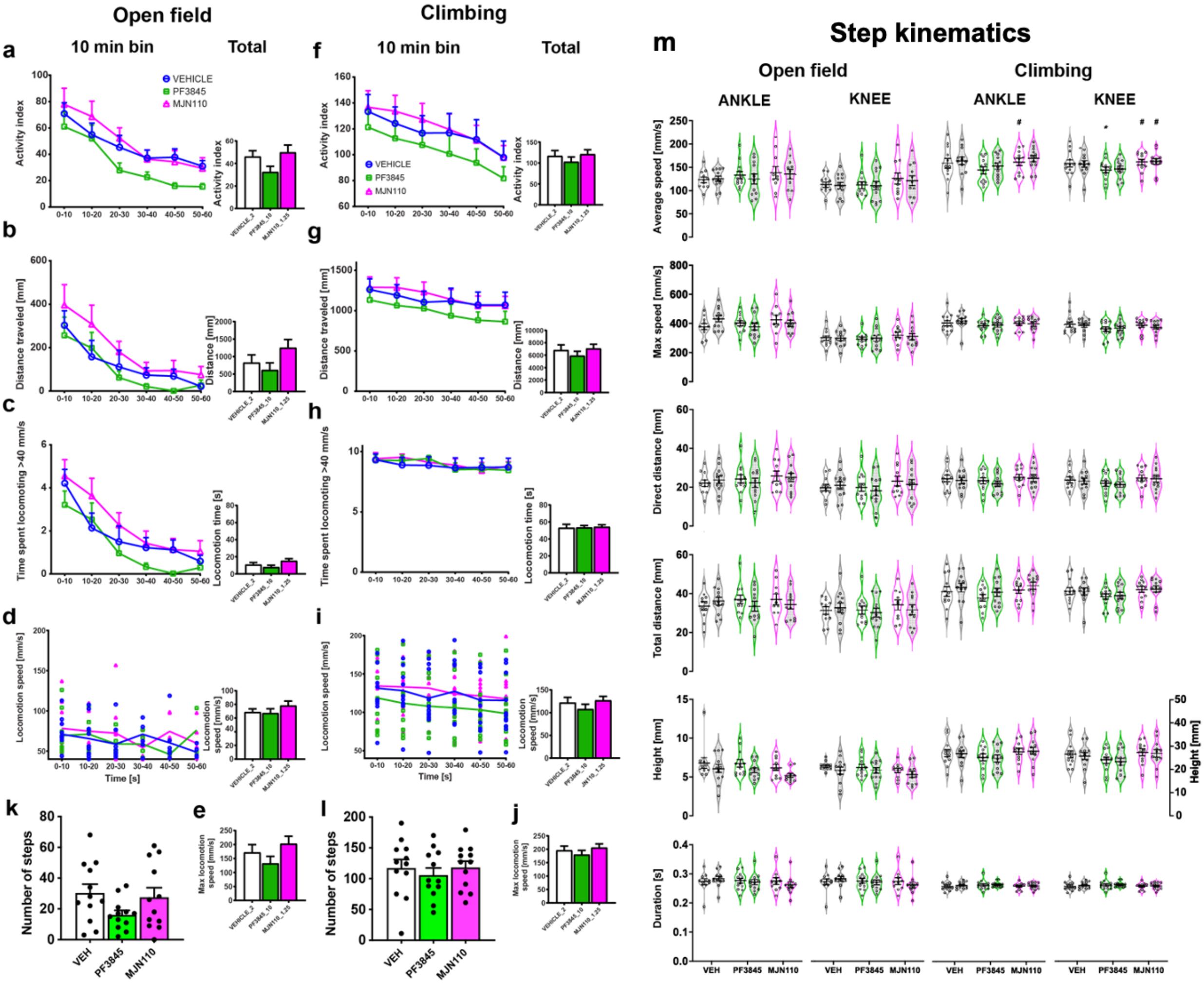


**Supplementary Figure 7. MAGL and FAAH inhibitors PF3845 (10 mg/kg) and MJN110 (1,25 mg/kg) produce small magnitude bidirectional effects on locomotor activity and step kinematics. a,** PF3845 slightly decreased activity index in the open field, but it did not reach statistical significance (p=0.052). **b,** Distance traveled **c,** time spent locomoting, were not significantly affected. **d,** No effect on average or **e,** maximum locomotion speed were observed in the open field. During climbing both inhibitors had similar less pronounced effects: **f,** activity index, and **g,** distance traveled, **h,** time locomoting, **l,** average and **j,** maximum speed were not significantly affected**. k,** Number of steps during the open field exploration was slightly decreased by PF3845 but it was not statistically significant (p=0.065). **l,** Number of steps during climbing was not affected. **m,** Small but significant effects on step kinematics were found on average swing speed of ankle (F (2, 22) = 3.91, p=0.35) and knee (F (2, 22) = 4.87, p=0.018) during climbing. *p<0.05, vs vehicle, #p<0.05 vs PF3845. Closed symbols denote significance in post hoc test vs vehicle. n=12 mice per group (randomized, counterbalanced within-subject design), mean ± SEM.

### Supplementary methods

#### Implementation and validation of the 3D Motion Capture system for rodents

Realization of marker-based 3D motion capture on small rodents poses several technical challenges that have been difficult to overcome [1,2]. First, skin-glued markers are of limited use in unrestricted rodents, as they attract the animal's attention and distort behavior. Moreover, markers made of soft materials will not survive the attention and require frequent replacements. While in restricted and strongly task-motivated animals (e.g. under food deprivation) such markers may survive sufficiently long for reliable tracking[3], our preliminary trials revealed that examining whole-body kinematics in freely-moving animals was impossible with skin-glued, plastic markers. To address this problem we established a method for permanent under-skin implantation of stainless-steel piercings at landmark locations, to which above-skin retroreflective markers are robustly attached (Fig. 1a, Suppl. Fig. 1 and 2, supplementary methods for details). After implantation of these markers at landmark locations, mice are habituated to them for two weeks prior to recordings.

Second, spurious reflections and distortions caused by any transparent surfaces between the camera and markers catastrophically degrade tracking performance that can only be compensated for by significant increase of tracking cameras. To allow reliable tracking with minimal number of cameras (6-7, placed at 60 degree angles around the recording arena in case of free horizontal exploration) we developed 2-week training protocol for the mice to learn not to leave the designated "arena" in the middle of the cameras despite absence of any physical barriers (Fig. 1a-c; Supp. Video 1). All animals used in the study learned this behavior without problems, allowing us to record their behavior not only on a horizontal, textured 30 x 30 cm surface (the "open field") but also several other settings. In this article we present the analysis of data of two tasks: a) exploration of the open field and b) vertical climbing behavior (Fig. 1c).

#### Piercing implantation procedure for motion capture in mice

Our preliminary studies indicated that implantation of stainless steel markers is the least invasive method of marker placement, and it will ensure natural behavior of the animal, as commonly-used plastic markers attract attention of animals, are difficult to attach (require strong glue causing skin irritation) but easy for animals to remove, which leads to chewing and swallowing. In contrast, stainless steel piercings do not cause any pain or discomfort and do not cause any change in the behavior. Another advantage of permanent markers is that animals can be tracked over time with the same localization of markers which enables powerful within-subject studies and high level of measurement precision.

#### Shaving and marking hole placement (10-20 min)

Male C57BL/6A mice (CLEA Japan) at 10-12 weeks of age and 20-25 g body mass at the time of implantation were used. Some distances should be adjusted when bigger or smaller animals are used.

After inducing anesthesia (2-3 %Isoflurane, Somnosuite, Kent Scientific, CT, USA) animals were shaven in places around areas where implants are located. Animals were placed evenly on the table making sure position of the animal is symmetrical and skin is distributed evenly. Hind legs were bent, and bottom of feet formed straight line perpendicular to the longitudinal axis of the mouse body (Suppl. Fig. 1). Skin was cleaned with 70% ethanol. Center between the hips and center between the shoulder blades was marked. Marked measuring stick (q-tip, prepared using precise calipers) showing appropriate distances between the holes to place the implant (e.g. 14 mm) was used to mark location of holes with blue marker (Suppl. Fig. 1). We used stainless steel barbel piercings with shaft length (distance between screw on balls) of 8 mm for the legs, 7 mm for hips and 6 mm for “coordinates“ and shoulder blades. In order to ensure durability of the implant, it is important to initially place it with double amount of the skin between two holes: e.g. 14 mm apart for 7 mm long piercing on the hips. If there is too little skin, the piercing will become loose and may fall out. Marks for holes for hip implant were made on top of the hips, symmetrically (14 mm wide both in shoulder blades and hips). 8-10 mm up from the hip center a dot was marked and another 2 symmetric dots were marked (14 mm wide) for the coordinate’s markers. It is important that they aren’t too close to hips because retroreflective markers are 2 mm larger that stainless steel marker. Marking place for upper leg hole was done by first moving each leg to maximally bent and half bent position and marking placement of the knee under the skin (according to how knee moves during walking), and hole was placed in between them. The lower leg hole was marked at 14 mm from the heel up on the center of the calves. The distance from the heel to the upper leg hole was around 28 mm. Symmetry of the marked dots on both legs was checked before implantation. It is important that lower leg holes are not too much to the front or back because they will slide behind the leg and will be invisible, they should be placed in the center of the calf. Ankle implants cannot be placed too low because retroreflective markers are larger and heavier and if they will fall too low with gravity when animal is standing up, markers will touch the floor and disturb animals walking.

**Placing implants (5-10 min)**

All piercings were disinfected with ethanol and dried before implantation. Forceps was used to lift the skin and and thick needle e.g. 18-21 G was used to break the skin at previously marked locations. Holes were then widened with super fine forceps (e.g. 55). The implants were placed in the holes and screwed on. There should be excess skin in between balls of the piercing but this will be adjusted during healing process. Animals were allowed 11-14 days for healing before starting recordings. During this time, we started behavioral training but we were avoiding exchanging implants before 11 days from implantation procedure because they are not fully healed at that time.

**Marker making method**

To achieve high retroreflectivity as well as durability sufficient for rodent experiments, we used 4 mm stainless steel barbel piercing screw on balls (Suppl. Fig. 1 and 2., da Phactory, Japan) covered with 3 layers: (1) retroreflective tape (3M, MN, USA), (2) polyethylene film (Trustco, Japan), and (3) UV hardened glue (Bondic, NY, USA) (Suppl. Fig 2.). Retroreflective tape was cut into narrow stripes (6 mm width) and then cut into slightly curved crescent shape in 1 mm width as seen in Suppl. Fig. 2. Each stripe is attached slightly overlapping on each other at the edges and the surface of a piercing is covered from the back to the front only in the coronal plane by this covering style. A round shape of uncovered metal surface in diameter of ~1-2 mm is covered by reflective tape as well cut into circular shape in diameter of ~2-4 mm. Small square pieces (2 cm X 2 cm) of polyethylene film are cut and tightly covered as a second layer on top of the piercings that are already covered by retroreflective tape. As the last step, liquid UV curable glue is applied and the third and outermost layer is formed by hardening with UV light. In order to save the sphericity of the markers, which is important for radial reflection, the marker should be continuously rotated during the hardening phase of the liquid UV glue. Gloves should be worn during the whole process to avoid fingerprints at the surface that would also result in distortion in reflectivity.

UV curable glue was chosen as the outermost layer of the piercing to protect the retroreflective tape, but the reflectivity of the tape is very sensitive to change in the refractive index of the medium and media other than air result in large distortion, and loss in reflected light. Therefore, polyethylene film coating is necessary before liquid UV glue application. Layer of thin plastic protects tape from contact with liquid that would diminish its retroreflective properties. Other methods including various retroreflective paints and sprays did not provide strong enough retroreflective signal on such a small volume and/or were not sufficiently resistant to damage. Number of materials were tested before identifying most successful marker design providing strong signal with excellent detection by the cameras, lightweight, and hard surface resistant to damage. All retroreflective markers are quality tested before they are used on animals.

**List of Tools & Materials:**

- Retroreflective tape, 3M Scotchlite 7610 Reflective Tape 1" Wide, Adhesive Backing (3M, MN, USA)
- Linear Low Density Polyethylene (LLDPE) film, TRUSCO sutorettifirumumini Micron 25 X W X/300 m, TSF2550 (Trustco, Japan)
- UV glue kit, Bondic, BD-SKCJ (Bondic, NY, USA)
- stainless steel piercing, straight stainless-steel barbel (da Phactory, Japan) 6-8 mm shaft, 3-4 mm ball
- Fixing material, Blu-Tack (Bostic Findley Ltd., Australia)
- Iris scissors, No:14095-11, (FST CA)
- Forceps, Dumont #5/45, No: 11251-35 (FST CA)
- Nitrile gloves
- UV light safety googles

### Behavioral training

After marker implantation, animals started 11-14 day training protocol to ensure proper wound healing and stable marker position. During those days animals were subjected to handling, acclimating, and task training in following order: 1) handling: a) taken out from the cage with the tube b) gently pulling by tail base on the outer part of hand or forearm c) scooping holding the tail gently d) scooping (2-3 days). 2) adaptation to arena: a) placement in circular 30 cm arena with 4 cm walls in brightly lit room (2-3 days) b) placement in circular area without walls and preventing mice from leaving the arena (taking back by the tail if 4 paws leave, tapping nose if 2 paws leave) (3-4 days). Mice are being kept in the arena for 2-5 min. 3) task training: a) vertical climbing on mesh b) horizontal climbing on string c) beam walking d) righting reflex (2-3 days, 1 min trial or less). Each animal is subjected to the same level of training. All animals are trained until they correctly perform task as expected. Markers that are worn by mice every day are being exchanged into retroreflective and larger markers 15 minutes before the start of recording. Mice are habituated to the process of manual exchange of markers that takes 3-5 min, so they do not need to be anesthetized, which could disturb the behavior. After finished recording markers are changed again and mouse is being returned to home cage.

### Calibrating motion capture recording for optimal tracking

We used the procedure provided by the hardware maker (Qualisys; https://docs.qualisys.com/getting-started/content/getting_started/running_your_qualisys_system/calibrating_your_system/calibrating_your_system.htm/) with slight modifications. First, to acquire uniform shapes of the markers while the animal moves in space, we focused the cameras at the outer perimeter of the behavioral arena instead of the center and used low aperture values (f = 2.8). This results in slight blurring of the small imperfections of the markers and more uniform image plane projection. Next, a L-shaped calibration bar and a T-shaped calibration wand are used according to the instructions provided to define the tracking volume so that X and Y directions are identical across all experiments. Great care was taken to ensure that the calibration volume was uniformly covered in the process. Calibration was considered acceptable when average triangulation residuals for all cameras were below 0.2 mm.

### Performance of the marker-based mouse motion capture system

A key question in assessing the potential and limitations of a novel recording method relates to the reliability and accuracy of the measurements. Here, we quantify them in terms of marker visibility (fraction of frames in a recording where a marker is visible or occluded) and location accuracy (how far is the recorded location from real position of a marker). Similarly to the approach taken in previous work[4], we calculated how many markers are occluded in each frame. As shown in Fig. 1d, left, vast majority of the frames had no markers missing and around 90% of frames had not more than one marker missing. Notably, the tracking quality was high even in climbing trials that were recorded with only 4 cameras.

When assessing the quality of tracking, the durations of occasional occlusions are of great significance as short gaps are trivial to be interpolated over while longer gaps require sophisticated approaches such as employment of machine learning and skeleton model fitting processing. As shown in Fig. 1d, right, most of the gaps terminated within 100 frames (corresponding to 0.3 s with 300 fps recording). Even for the most challenging ankle markers that by necessity are fully occluded when the animal is in an idle, inactive position, the occlusions were on average resolved within 50 frames (0.15 s). We also could confirm that there were no differences in marker tracking performance for left and right sides of the animals, evidencing there were no biases in how the cameras were positioned with respect to the behavioral arena configuration (Student's paired t-test for left and right legs for 30 1-minute-long recordings, p = 0.7).

The accuracy of triangulation-based 3D tracking depends critically on the uniformity of marker shape and its projection into the camera image plane. To assess the performance of our system in this respect, we took advantage of the fact that the visible marker pairs reside at the ends of metal rods of fixed and known length, and the deviation of a marker's tracked position relative to it's pair can be considered as its error. Fig. 1e shows that the distance between paired markers varied on average no more than 0.5 mm even for the body parts most occluded by hair (shoulder blades); notably, assuming independent errors for markers, leg marker distance deviations of around 0.2 mm indicates that a tracked position of a single marker could be expected to be within 100 micrometers of its real position.

For step detection (Fig. 1f,g), we considered that a swing consists of a period of high-velocity ankle movement bordered with rapid acceleration and deceleration events. Fig. 1g depicts schematically how speed (red trace) and acceleration (green trace) relate to the vertical position of the ankle (blue trace) and determine our "swing" segment. To validate this approach and define its limitations, we compared the number of steps detected with those identified in recordings by an experienced observer (BIJ). As shown in Fig. 1h (left), in 30 60-s trials mice took on average 92.9 steps, out of which 62.1 steps were detected by the algorithm. Taking a step further in detailing the reliability, we chose 3 animals representing the animals with average, highest (best) and lowest (worst) percentage of steps automatically detected and determined the number of missed steps and false positives in these samples (Fig. 1h, right, Suppl. Fig. 3). In this representative set of tracked steps, on average 81 out of 108 steps were correctly detected; importantly, false positives were below 1%, attesting to the validity of our step-detection principle.

Notably, post-processing of the trajectories with machine learning or model fitting approaches could be used to further "deconvolve" the tracking data. However, we chose to avoid unnecessary complexity in the analysis as the performance was sufficient for simultaneous tracking of animal bodies in various behavioral contexts and detecting a representative sample of step trajectories.

### Source code and kinematic feature extraction

Kinematic parameters were extracted by custom made code in Matlab (2019a) run on MacOS Monterey 12.1. All code will be made available at github repository at the time of publication and link will be provided here.

Key parameters computed from the 3D trajectories:

- mouse speed: displacement of mean hip markers in 1 second
- locomotory episodes: frames in which mouse speed remained over 40 mm/s for longer than 100 frames. The threshold value was determined based on observations of 10 randomly selected videos from the data pool.
- locomotory bouts: blocks with peaks in speed with minimal prominence of 15 mm/s, and the nearest matching local minima.
- Activity index: mean of all markers' speed (smoothed over 10 frames).
- Hip-shoulder height: distance of mean hip and shoulder markers in the dimension orthogonal to locomotory plane (horizontal or vertical for open field and climbing, respectively).
- Hip-shoulder distance: euclidean distance of the mean hip and shoulder markers.
- Step detection: Swing segments of steps were delineated based on kinematic rather than positional features of ankles, understanding that limb joints do not simply move up and down during stepping. The underlying principle is that during swings, ankles move (in any direction) with a high speed relative to the ground (either floor or the wheel), and a fully-resolved swing must be bordered with acceleration and deceleration events, reflecting take-off and touch-down events. From detected high-speed ankle events, we excluded those for which we could not determine precise acceleration and deceleration peaks. While this approach necessarily omits steps with "creeping" take-off or touch-down events (that could

be more prominent in the moments leading to ceasing locomotion), we chose it over more lenient inclusion criteria in order to reduce the probability of falsely-identified steps (such as those related to scratching or postural shifts in place) from contaminating the measurements.

- Step measurements (calculated for both left and right hind limb separately):
  - swing height: maximal displacement of the ankle marker during a swing, in dimension orthogonal to the locomotory plane (horizontal or vertical)
  - swing direct distance: euclidean distance of the ankle or knee marker position at frames defined as swing start and swing end;
  - swing total distance: length of the 3D trajectory of the ankle or knee marker between the frames defined as swing start and swing end;
  - swing mean and max speed were calculated from the segments designated as swings, without smoothing.
  - stance width: sum of the mean distance of left and right ankles to the centerpoint of hip and coordinate markers
